# The combination of elevated neuronal activity and mitochondrial damage induces Pink1-dependent mitophagy in axons

**DOI:** 10.64898/2025.12.26.696432

**Authors:** Serena R. Wisner, Chris Stein, Erin E. Smith, Mrinalini Hoon, Catherine M. Drerup

## Abstract

Mitochondria are critical for synaptic function. At the synapse, mitochondria produce ATP and buffer calcium, both of which are required for synapse function. Defects in mitochondrial maintenance are linked to neurodegenerative disease, yet we know little about what regulates the need for mitophagy at the synapse. We assessed the impact of neuron type, activity, and mitochondrial damage on mitophagy rate in axons of larval zebrafish. Using electron and confocal microscopy, we show that mitophagy occurs in the axon terminal of postsynaptic sensory neurons and presynaptic motor neurons at similar rates. Increasing neuronal activity or mitochondria damage does not impact the amount of mitophagy in axons. Only by combining neuronal activity and mitochondrial damage does the rate of mitophagy increase in the axon and this increase requires Pink1. Together, our data support a model in which increased mitophagic demand in axons is rare and uniquely sensitive to Pink1 disruption.

## Introduction

Mitochondrial function is critical for neuronal health and function. At the synapse, mitochondria provide ATP and buffer calcium to support synaptic vesicle recruitment and recycling^1–3^. To support viability and function of a long-lived cell like a neuron, mitochondrial populations in the cell must be maintained. This is particularly critical in the axon, an area of high metabolic activity^4^. One critical process for mitochondrial population maintenance is the selective degradation of damaged mitochondrial components through the process of mitophagy^5^.

Perhaps not surprisingly, mutations in key mitophagy genes have been linked to neurodegenerative diseases. Most notably, mutations in genes encoding Pink1 and Parkin cause Parkinson’s disease^6–10^. In Pink1/Parkin-dependent mitophagy, Pink1 acts as a mitochondria damage sensor, stabilizing on the outer mitochondria membrane (OMM) after mitochondria matrix depolarization^9^. This leads to the phosphorylation of downstream targets, Parkin activation, and Parkin-dependent ubiquitination of the damaged mitochondria^10–12^. Interestingly, mitophagy can occur in the absence of Pink1, suggesting context-specific activation of this pathway^13,14^. Given the crucial link between mitochondrial turnover and neurodegeneration, understanding the physiology which drives the need for Pink1-dependent mitophagy in neurons is needed.

One factor which may influence risk of neurodegeneration with disrupted mitophagy is neuron type. This is most clearly shown in Parkinson’s disease in which mutations in Pink1 result in death of dopaminergic neurons of the substantia nigra (SNc)^15^. Other neuron populations are largely unaffected, particularly in early stages of the disease. The reason for this is largely unknown, but it has been postulated that increased reliance on mitochondrial function and high levels of synaptic activity may underlie sensitivity. This is supported by data comparing dopaminergic neurons of the SNc with those of the ventral tegmental area (VTA). SNc dopaminergic neurons have higher levels of oxidative phosphorylation (OXPHOS) and an associated increase in oxidative damage compared to those in the VTA. Furthermore, VTA neurons are smaller and have fewer synapses than SNc dopaminergic neurons^16^. Together, this points to a profound difference in risk due to neuronal activity and mitochondrial function. However, how neuronal type, mitochondrial damage, and neuronal activity impact mitophagy rate has not been specifically addressed.

We used live imaging in larval zebrafish to determine how neuron type, activity, and accrued mitochondrial damage impact mitophagy rates in the axons of larval zebrafish. Specifically, we focused on presynaptic primary motor neurons (pMN) and sensory axons of the posterior lateral line (pLL) system in 5 day post-fertilization (dpf) larvae. At this stage, primary larval nervous system development is complete, and animals must interact with their environment using active sensory and motor circuits for survival. Using neuronal populations, we determined the basal rate of mitophagy in the axon, near the synapse, is almost identical between these very different axons. Using behavior-driven increases in neuronal activity and genetic induction of mitochondria damage, we show that individually these stressors do not change axonal mitophagy rate on their own. Rather, when combined, these factors increase the demand for mitophagy in the axon, near the synapse. This increase in mitophagy demand in the axon requires Pink1 function. Together, our results demonstrate the combinatorial relationship between physiological factors regulating mitophagy. Furthermore, based on our data, mitophagy in the axon does not require Pink1 pathway unless mitophagy drive is increased. This points to a model of context specific Pink1-dependent mitophagy which, when disrupted, may predispose some neuronal populations to neurodegeneration.

## Results

### Mitochondrial derived vesicles (MDVs) are the primary method of mitophagy under basal conditions

Mitochondrial degradation can occur through two primary paths: 1) damaged mitochondrial components can be packaged into MDVs which are subsequently degraded; or 2) the whole mitochondrial organelle can be engulfed and degraded. The primary means of mitophagy for maintenance in axons has not been addressed. We analyzed mitochondrial budding events, a precursor to MDVs, and whole organelle mitophagy in serial block face scanning electron microscopy (SBFSEM) of the larval zebrafish posterior lateral line (pLL) system afferent and efferent axons. In the pLL system, afferent axons extend from the pLL ganglion situated behind the ear while efferent axons extend from the hindbrain into the periphery to innervate sensory organs throughout the trunk^17^. These sensory organs contain hair cells which respond to water movement. Mechanical deflection of the hair cell stereocilia by water movement results in glutamate release from ribbon synapses situated at the base of the cell onto post-synaptic afferent pLL axon terminals (Figure 1A,B)^18,19^. Hair cell activity is modulated by presynaptic efferent axons that provide inhibitory feedback (Figure 1A,B)^20^. Afferent and efferent axons offer a unique opportunity to assess pre- (efferent) vs post- (afferent) synaptic axon mitochondrial parameters in the same sensory organ. First, we assessed mitochondrial size and found that, similar to previous studies in other systems^21,22^, efferent (presynaptic) axon terminal mitochondrial volume was significantly lower than afferent (postsynaptic) axon terminal mitochondria (Figure 1C, Log2 scale, afferent: 1.369 ± 0.156 vs. efferent 0.378 ± 0.209, p=0.0003, mean ± SEM).

**Figure 1.**
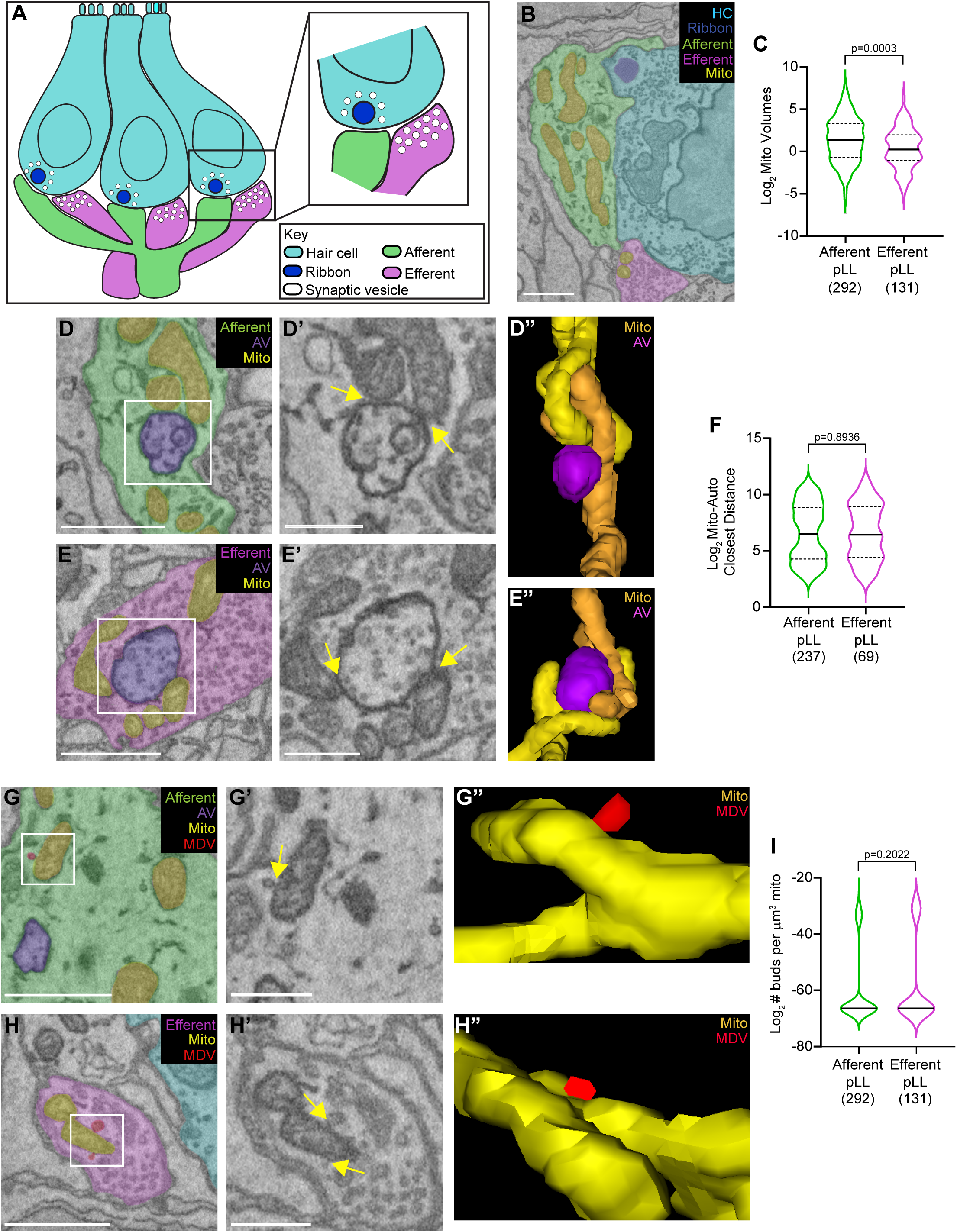
Occurrence of mitochondria budding events across cell types. **(A)** Schematic of the zebrafish posterior lateral line neuromast. Hair cells (cyan) are activated by water movement. This triggers the release of neurotransmitter at ribbon synapses (dark blue) at the hair cell base. Afferent axon terminals (green) contain the post-synapse. Efferent axon terminals (magenta) provide modulatory feedback onto the hair cell. (**B**) SBFSEM image of hair cell (cyan) with a ribbon synapse (dark blue) contacting an afferent axon terminal (green) and receiving input from the efferent axon terminal (magenta). Mitochondria (yellow) can be seen in both axon terminals. (**C**) Quantification of mitochondria volumes in afferent and efferent axon terminals (Log2 transformed; Mean ± SEM, afferent: 1.369 ± 0.156 vs. efferent: 0.378 ± 0.209, p=0.0003, ANOVA). (**D, E**) Image of an autophagic vesicle (purple) contacting a mitochondria (yellow) in afferent (**D**, green) or efferent (**E**, magenta) axon terminals. White box indicates region enlarged in **D’, E’** with 3D model shown in **D”** and **E”** for the afferent and efferent axon terminal respectively. Arrows indicate contact site between mitochondria and the autophagic vesicle. (**F**) Quantification of the closest mitochondria-autophagic vesicle distance (Log2 transformed; Mean ± SEM, afferent: 6.579 ± 0.164 vs. efferent: 6.613 ± 0.297, p=0.8936, Wilcoxon). (**G, H**) Images of an afferent (**G**) and efferent (**H**) mitochondria (yellow) with a mitochondria budding event (MDV, red). White box indicates region enlarged in **G’**, **H’** with 3D model shown in **G”** and **H”** for the afferent and efferent axon terminal respectively. Arrows indicate site of budding event.(**I**) Quantification of the number of buds per um^3^ of mitochondria in afferent and efferent axon terminals (Log2 transformation; Mean ± SEM, afferent: -60.123 ± 0.81 vs. efferent: -58.272 ± 1.20, p=0.2022, ANOVA). For all, sample size below graph is number of individual mitochondria from 3 neuromasts from 3 SBFSEM datasets.

Next, we assessed mitochondrial budding events and whole mitochondrial engulfment in our SBFSEM datasets. While we observed close contacts between mitochondria and autophagic vesicles in both types of axons (Figure 1D-F), we failed to find any examples of engulfed mitochondria in our dataset (9 afferent axon terminals, 4 efferent axon terminals from 3 sensory organs). Conversely, mitochondrial budding events to produce MDVs were readily apparent. Budding events, characterized by the presence of an outcropping from a mitochondria with a neck constriction (indicating bud fission), were present in 19% and 23% of afferent and efferent mitochondria respectively (Figure 1 G,H). Additionally, when analyzing number of buds per mitochondrial volume, we found no difference in the number of budding events between neuron types (Figure 1I). The size of the mitochondrial budding events were the same in pLL axon types as well (afferent: 58.9 nm ± 2.63, efferent: 53.9 nm ± 3.00), which are consistent with previous findings of MDV size in electron micrographs^23^. Together, our data indicate that MDV production may be the predominant means for mitophagy in both afferent and efferent axon terminals.

### Mitophagy rates are unaffected by neuronal type

The consistent frequency of mitochondrial budding in our SBFSEM datasets suggests that basal mitophagy rate may be tightly controlled in axons. However, mitochondrial budding events are also linked to mitochondrial biogenesis. To delineate mitophagy verses mitochondrial biogenesis, we next assessed mitophagy using live imaging of a mitophagy reporter. For our subsequent work, we chose to compare presynaptic primary motor neurons (pMNs) and postsynaptic pLL sensory axons in larval zebrafish (Figure 2A-B). Both neuron types are superficially located, with stereotypical branching patterns for ease of identification^17,24^. However, motor but not sensory neurons are highly susceptible to neurodegeneration in disease states across model species and in humans^25–27^. To examine the rate of mitophagy in pLL and pMN neurons, we used a dual tagged GFP-mCherry indicator localized to the mitochondria matrix (Figure 2C)^28,29^. GFP is quenched in acidic environments, such as that found in an autolysosome, allowing quantification of mitochondrial area undergoing mitophagy (mCherry only) relative to the total mitochondrial population (GFP and mCherry positive).

**Figure 2.**
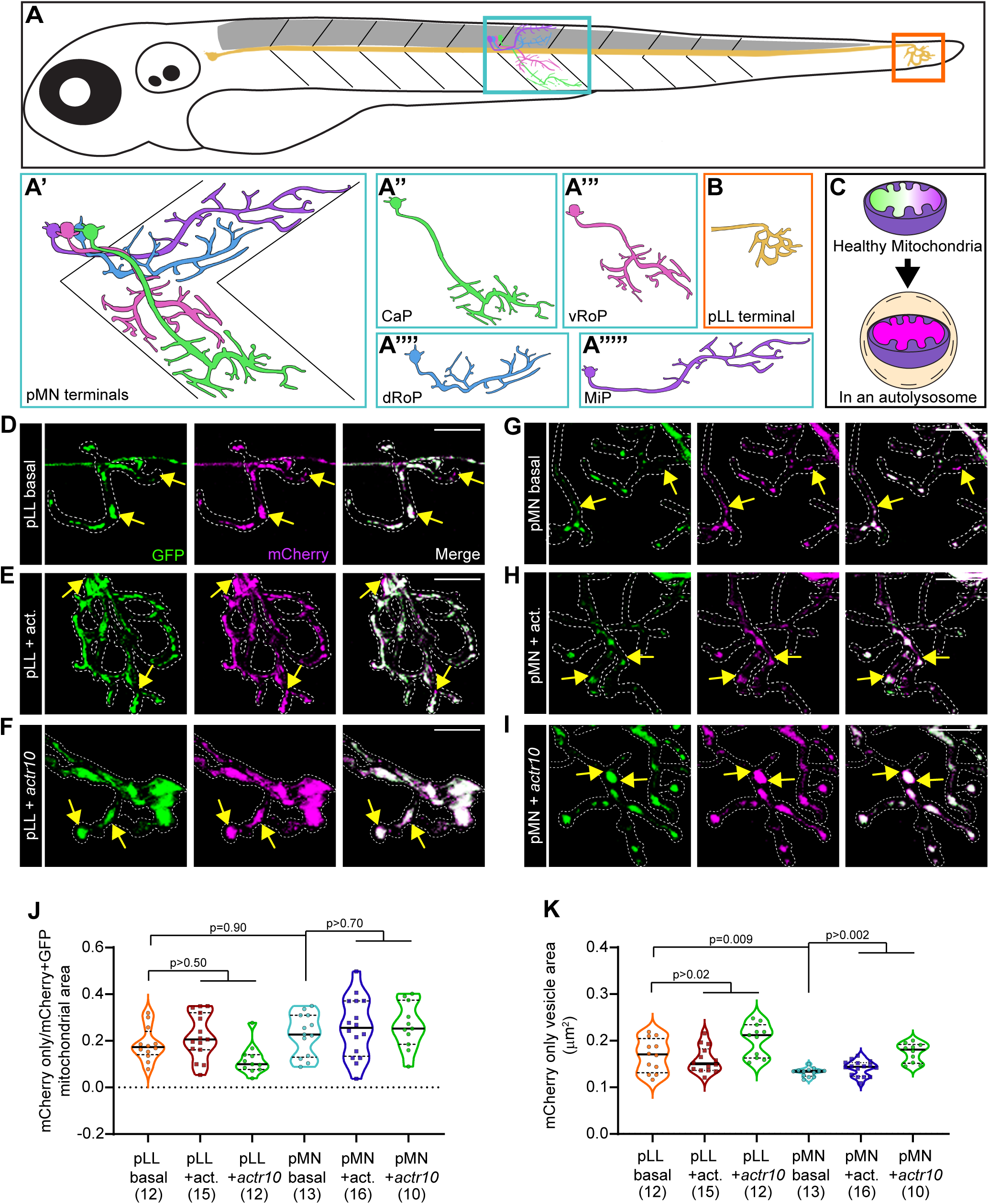
Neuronal type, activity and damage do not independently alter axonal mitophagy. (**A**) Schematic of the larval zebrafish. Sensory and motor neurons schematized. The posterior lateral line (pLL) axon is shown in yellow with the axon terminal boxed in orange. Primary motor neurons (pMNs) are boxed in blue. Spinal cord is shown in grey. **(A’-A””’)** Illustrated examples of distinct primary motor neuron innervation patterns; Caudal primary (CaP) (**A’’**), ventral rostral primary (vRoP) (**A’’’**), dorsal rostral primary (dRoP) (**A’’’’**), and middle primary (MiP) (**A’’’’’**). (**B**) Illustrated example of the pLL axon terminal. (**C**) Schematic of the mitophagy indicator. In healthy mitochondria, both GFP and mCherry (magenta) fluoresce, giving a white appearance. In an acidic environment, such as an autolysosome, the GFP signal is quenched, leaving an mCherry only signal. (**D-F**) Representative images of mitophagy in wild type (WT) pLL axon terminals without (**D**) and with (**E**) increased neuronal activity (+act.) and in an *actr10* mutant (**F**). Arrows indicate portions of mitochondria undergoing mitophagy (mCherry only). (**G-I**) Representative images of mitophagy in a portion of a wild type pMN axon terminal without (**G**) and with (**H**) increased neuronal activity or in an *actr10* mutant (**I**). Arrows indicate portions of mitochondria undergoing mitophagy (mCherry only). (**J**) Quantification of the proportion of mitophagic area (mCherry only) to total mitochondria area (mCherry+eGFP positive; ANOVA with Tukey HSD post-hoc contrasts). (**K**) Quantification of the average mCherry only mitochondrial area (Steel-Dwaas all pairs). Sample size, indicated below graph, represents individual larvae. Scale bar = 10 μm.

We expressed the mitophagy indicator using zygotic microinjection of plasmid DNA and quantified the relative area of acidified mitochondria in pLL and pMN axon terminals. First, we confirmed efficacy of this mitophagy indicator in our neurons using live imaging. We expressed the mitophagy indicator with the lysosome associated protein Lamp1 tagged with TagBFP2 and assessed colocalization. mCherry only organelles strongly colocalized with Lamp1-positive lysosomes (Figure S1A). Then, we assessed the basal rate of mitophagy in sensory and motor neuron axons using the mitophagy indicator. This demonstrated a consistent proportion of mitochondrial area undergoing mitophagy in both axonal types, around 20% (Figure 2D,G,J; pLL: 18.6% ± 2.1; pMN: 22.6% ± 2.5). Quantification of mCherry only mitochondrial size estimates that the portions of mitochondrial undergoing degradation are approximately 0.100-0.200 μm^2^, consistent with MDV-dependent mitophagy (Figure 2K; pLL: 0.168 ± 0.01; pMN: 0.132 ± 0.002)^30^. While larger than the mitochondrial budding events we observed with SBFSEM, this is likely a size overestimate due to the limit of optical resolution of confocal microscopy (∼300 nm). Together, our electron microscopy and light level analysis demonstrate consistent levels of mitophagy in pMN and pLL sensory axons and suggests that MDVs are the primary mechanism of mitochondrial maintenance in this neuronal compartment.

### Neuronal activity and accrued mitochondrial damage on their own do not change rate of mitophagy

Because neuron type alone did not change the requirement for mitophagy, we next asked if increased neuronal activity could influence the rate of mitophagy. To activate pLL sensory axons, we used water flow. As discussed above, this afferent neuron is stimulated in response to water movement around the larvae. We chronically stimulated the pLL circuit by rocking larvae at 85 rpm for 36 hrs and tested mitophagy as described above. pLL stimulation did not change the proportion or size of mitochondria undergoing mitophagy (Figure 2D,E,J,K). Then, we stimulated pMNs to assess if increased activity increased rate of mitophagy. To stimulate pMNs, we incubated larvae in 100 μM NMDA for 20 hrs. NMDA treatment has previously been used by us and others to increase swimming frequency and the synaptic firing rate at pMN axon terminals^31–34^. To confirm treatment efficacy, we used antibody labeling for phosphorylated ERK (pERK) and demonstrated an increase in pERK labeling in the ventral spinal cord after NMDA treatment (Figure S1B-D)^35^. The proportion of mitochondrial area undergoing mitophagy and size of mitophagic vesicles were not changed by chronic increases in pMN activity (Figure 2G,H,J,K). Together, these data demonstrate that mitophagic rates are highly consistent across these disparate neuronal types and are not altered by behaviorally-associated increases in axonal activity.

In cultured cells, drugs used to disrupt mitochondrial membrane potential or impair the electron transport chain cause a rapid and robust increase in mitophagy across the mitochondrial population^36–38^. Rather than immediately disrupt the entire mitochondrial network, we wanted to selectively disrupt mitochondrial health in axons by causing accrued mitochondrial damage, akin to age-related neurodegenerative disease. To do this, we used a mutant zebrafish line (*actr10^nl^*^15^; *actr10* hereafter) that we have previously shown has defects in pLL and pMN axon mitochondrial health and function^31^. In *actr10* mutants, mitochondria cannot be removed from the axon terminal via dynein-dependent transport, leading to accumulation of mitochondria in the distal axon^39^. In *actr10* mutant pLL sensory axons, axon terminal mitochondrial health measures are reduced including lowered mitochondrial matrix potential, decreased calcium buffering, and increased reactive oxygen species accumulation^31^. Similarly, pMNs in *actr10* mutants display loss of ATP production but other parameters of mitochondrial health have not been tested. We first assessed mitochondrial matrix potential in pMNs using the cationic dye tetramethylrhodamine ethyl ester (TMRE) to determine if they could be used as a model of axonal mitochondrial damage. TMRE accumulates in the negatively charged mitochondrial matrix so level of fluorescence can be used to assess relative mitochondrial health. Similar to pLL axons, pMN axons display lower mitochondrial matrix potential in *actr10* mutants, confirming these organelles are damaged (Figure S1E-G)^31^. Then we assessed mitophagy in *actr10* pLL and pMN axons. Surprisingly, accrued mitochondrial damage in this line does not significantly increase the proportion of mitochondria undergoing mitophagy (Figure 2D,F,G,I,J). Interestingly, there is a slight increase in size of the mitophagic area which could suggest an increase in whole organelle mitophagy in both pLL and pMNs (Figure 2K). However, it does not drive up the overall proportion of mitochondrial undergoing mitophagy in these axons. Together, our data demonstrates that basal mitophagy is tightly regulated to maintain population health, likely through MDV production, and that physiologically relevant levels of increased neuronal activity and mitochondria damage do not drive a significant increase in mitophagy in axons.

### Neuronal activity and mitochondrial damage together increase mitophagy rates in axons

Neurodegenerative disease risk has been linked to neurons with high levels of synaptic activity and associated accrued mitochondrial damage^16,40^. Based on this, we next asked if a “two-hit” model, i.e. high activity and accrued mitochondrial damage, was necessary to drive an increase in the rate of mitophagy, suggesting increased need for this cellular process. To test this, we expressed the mitophagy indicator in wildtype and *actr10* pMNs, stimulated them with 100 μM NMDA to increase swimming, and assessed mitophagic rate. We focused on pMNs for this analysis as motor neurons are more sensitive than sensory neurons to degenerative disease states. When we quantified the proportion of mitophagic area in pMN axons, we found a significant increase in axonal mitophagy in *actr10* pMNs after neuronal stimulation (Figure 3A,B,D). We confirmed

**Figure 3.**
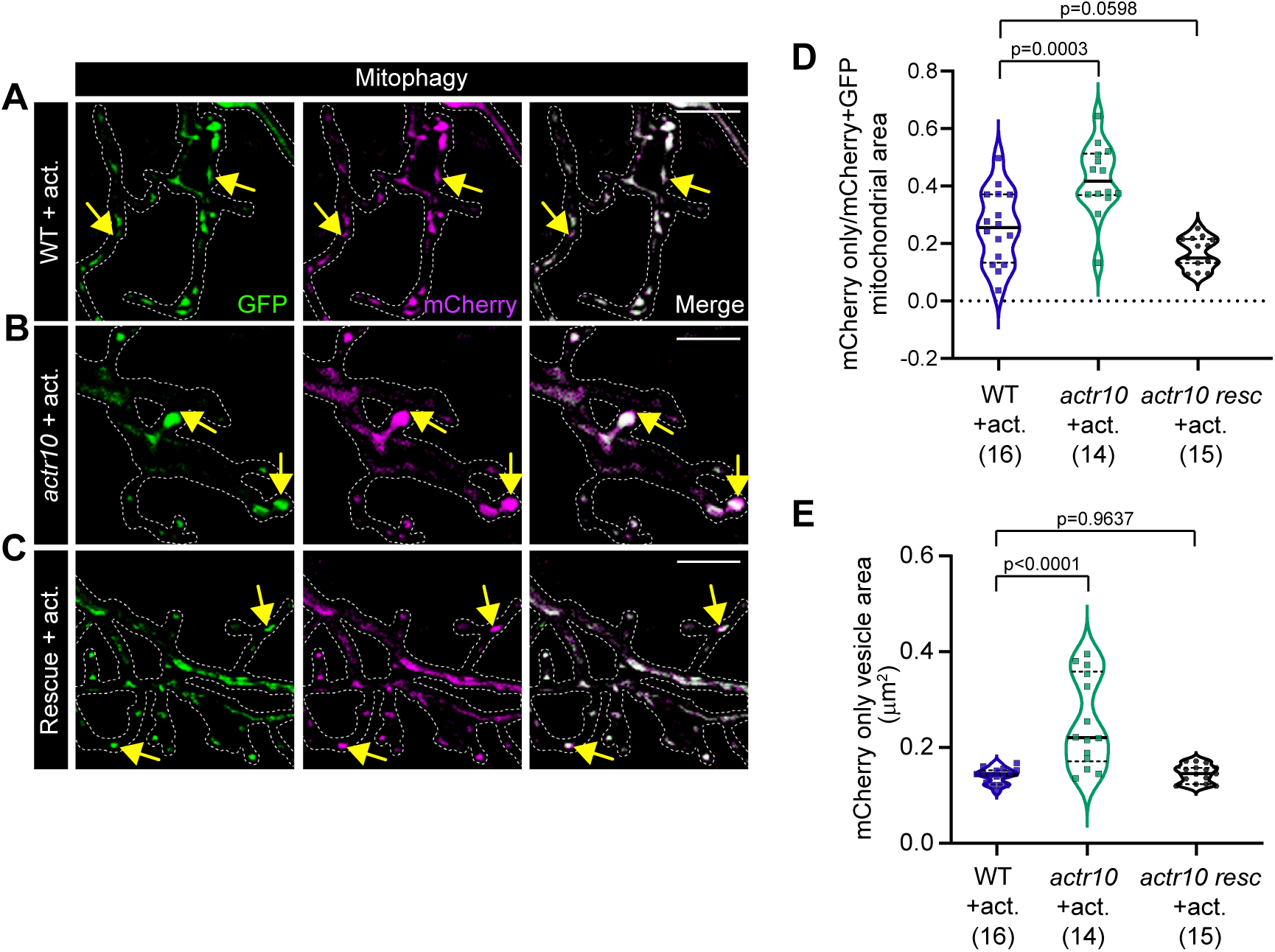
Mitochondria damage and neuronal activity synergistically increase axonal mitophagy. (**A**-**C**) Representative images of portions of pMN axon terminals labeled with the mitophagy indicator in wild type (WT; **A**), *actr10* mutants (**B**), and *actr10* mutants with neuronal rescue of wild type Actr10 (Rescue; **C**) with increased neuronal activity. Arrows indicate portions of mitochondria undergoing mitophagy. Scale bar = 10 μm. (**D**) Quantification of the proportion of mitophagic area (mCherry only) to total mitochondria area (Steel-Dwaas all pairs). (**E**) Quantification of the average mCherry only mitochondrial area (Steel-Dwaas all pairs). Sample size, indicated below graph, represents individual larvae. increased mitophagy was at due to loss of Actr10 in highly active pMNs using rescue analysis. We expressed wild type Actr10 in pMNs using a stable transgenic line (*actr10^nl22TG^;* RFP-Actr10 hereafter^31^), stimulated with NMDA, and assessed mitophagy. RFP-Actr10 expression in neurons suppresses increased mitophagy (Figure 3A-D; wild type + stimulation: 25.6% ± 3.2, vs. *actr10* + stimulation: 42.3% ± 1.4, vs. *actr10* rescue + stimulation: 11.8% ± 1.9). Quantification of size of mitophagic vesicles also demonstrates an increase in *actr10* mutants after stimulation that is rescued in the RFP-Actr10 transgenic line (Figure 3E, wild type + stimulation: 0.141 ± 0.002, vs *actr10* + stimulation: 0.253 ± 0.025, vs *actr10* rescue + stimulation: 0.146 ± 0.005). Together, our results show that accrued mitochondria damage and high neuronal activity can drive increased mitophagy in pMN axons. The increased size of mitophagic vesicles again suggests there may be a shift to either larger MDVs or whole organelle mitophagy with this two-hit model of neuronal mitochondrial damage.

Increased mitophagy requires a related increase in overall measures of autophagy. We measured autophagy using expression of a dual-tag mCherry-GFP-LC3 reporter in motor neurons of zebrafish larvae (Figure S2A)^41–44^. As with the mitophagy reporter described above, GFP is quenched in the acidic environment of an autolysosome after autophagosome-lysosome fusion. The mCherry only positive autophagosome area is therefore a readout of active autophagy in axons. We expressed this indicator in wild type and *actr10* mutant pMNs with and without NMDA-mediated stimulation and assessed the proportion of mCherry only autophagosomes (Figure S2B-E). While there is a trend towards higher axonal autophagy in *actr10* mutants, adding increased neuronal activity makes this group significantly elevated compared to wild type animals (Figure S2F, wild type basal: 18.1% ± 2.5, vs. wild type + stimulation: 22.4% ± 1.7, vs *actr10* basal: 26.7% ± 2.59, vs. *actr10* + stimulation: 29.6% ± 2.7). The increase in pMN axon autophagy caused by the combination of mitochondrial damage and neuronal activity is less than the increase in mitophagy (autophagy increase: 24%; mitophagy increase: 39%) suggesting mitochondrial components are the primary target of autophagy which is increased in the axon in this two-hit condition. Together, our data demonstrates that accrued mitochondrial damage in a highly active neuron increases mitophagy demand.

### Pink1-mediated mitophagy is required to meet increased mitophagy demand

Several mechanisms have been described which target damaged mitochondria for mitophagy. Pink1-mediated mitophagy is the most well studied mitophagy pathway which, when disrupted, leads to early onset Parkinson’s disease^9,10,45,46^. Pink1-dependent mitophagy is initiated when mitochondrial matrix polarization fails, like we see in *actr10* mutant pMN axons (see Figure S1). Based on this, we asked if Pink1-dependent mitophagy was responsible for the increase in axonal mitophagy observed after accrued mitochondrial damage with increased neuronal activity. To address this, we used CRISPR/Cas9-mediated mutagenesis to create a frame-shift mutation in the first exon of *pink1* in zebrafish (*pink1^uwd^*^13^, hereafter, *pink1,* Figure S3A). Large deletions oftentimes cause nonsense-mediated mRNA decay, indicative of loss of gene function. Indeed, in situ hybridization demonstrated that *pink1* mRNA is lost in our *pink1* mutant line (Figure 4A-C; wild type: 1 ± 0.099, vs *pink1*: 0.356 ± 0.048) confirming this is a loss of function mutation. To determine if Pink1 is required to increase mitophagy in response to mitochondrial damage and neuronal activity, we assessed mitophagy in *actr10;pink1* double mutants. Compared to wild type, *actr10* mutants again show an increase in pMN axonal mitophagy in response to neuronal activity; however, the *actr10;pink1* double mutants fail to upregulate pMN axonal mitophagy (Figure 4D-G; wild type + activity: 22.6% ± 1.6; *actr10* + activity: 33% ± 1.8; *actr10;pink1* + activity: 22.6% ± 1.7). Additionally, we noted a decrease in the number of primary axonal branches and reduction in total axonal area in the double mutant line, which could indicate axon degeneration (Figure S3B-G). Together, our data support a model in which accrued mitochondrial damage and sustained neuronal activity increase Pink1-dependent neuronal mitophagy in the axon. Loss of Pink1 impairs mitophagy upregulation which likely contributes to risk of neurodegeneration.

**Figure 4.**
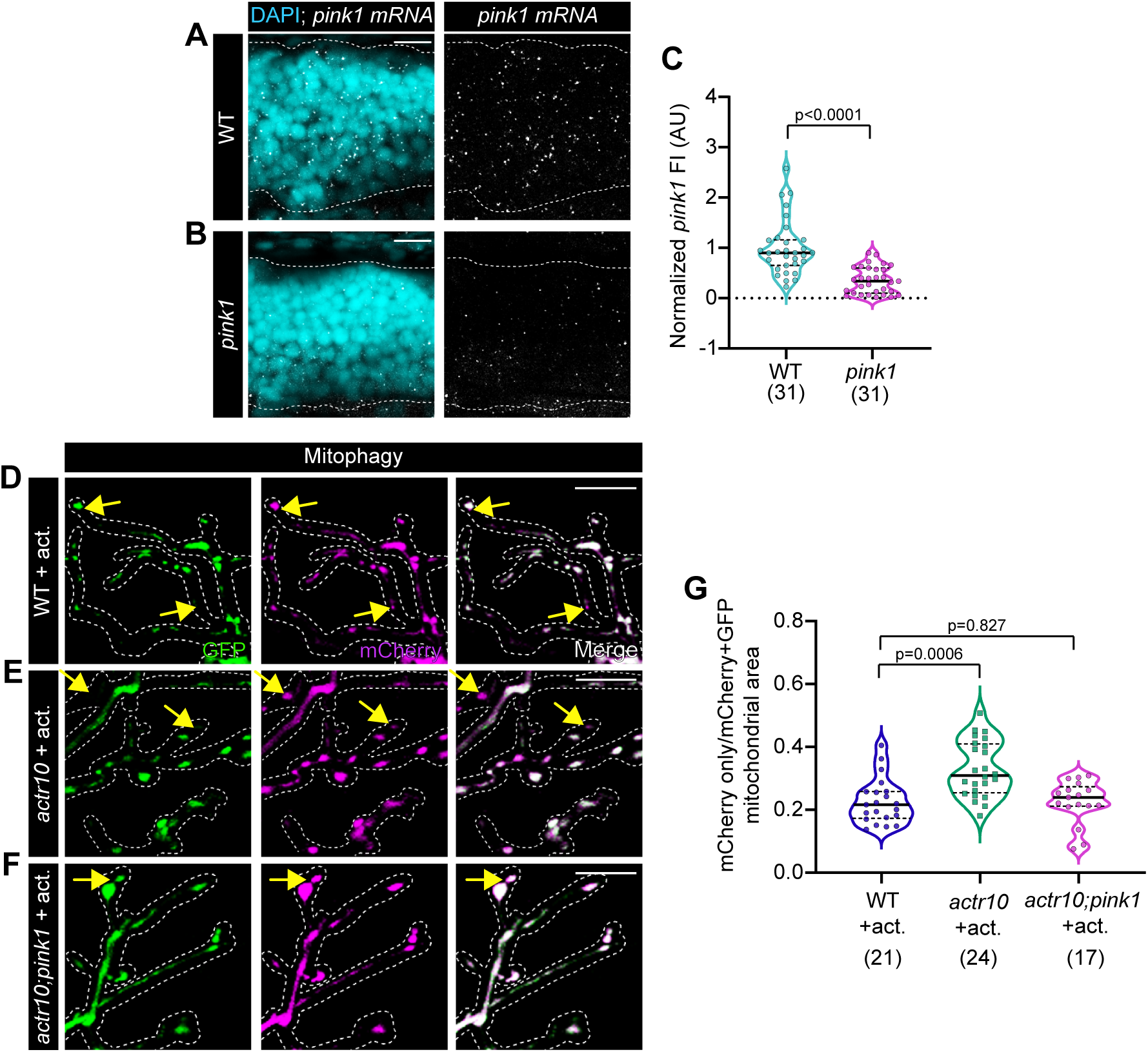
Increased mitophagy requires Pink1. (**A**-**B**) Representative images of *pink1* HCR RNA FISH (white) in spinal cord (DAPI – cyan) of wild type (WT; **A**) and *pink1* mutant (**B**) zebrafish larvae. (**C**) Quantification of normalized *pink1* mRNA fluorescence intensity (FI; data normalized to wild type mean; Wilcoxon). (**D**-**F**) Representative images of pMN terminals of wild type (**D**), *actr10* mutants (**E**), and *actr10;pink1* double mutants (**F**) with increased neuronal activity expressing the mitophagy indicator. Arrows indicate mCherry only mitochondria. (**G**) Quantification of proportion of mCherry only mitochondrial area relative to total mitochondria area (Steel-Dwaas all pairs). Scale bars = 10 μm. Sample size, indicated below graphs, represents individual larvae.

## Discussion

Despite its importance for neuronal survival and function, an understanding of the driving factors that stimulate the need for mitophagy and the mechanism of mitophagy upregulation in the axon is limited. Here, we addressed the impact of neuronal type, neuron activity, and mitochondrial damage on the rate of mitophagy in zebrafish neurons. We found no difference in the basal mitophagy rate between sensory and motor neurons. Neither increasing axonal mitochondrial damage nor neuronal activity changed mitophagy rate on their own. Rather, it was the combinatorial interaction of activity and mitochondrial damage that led to an increase in mitophagy in the axon.

Mitophagy upregulation in this “two-hit” model required Pink1-mediated mitophagy. Together, our work builds towards a model of mitophagic modulation in neurons in which mitophagy rate is largely hardwired. Increasing the demand for Pink1-dependent mitophagy requires accrued mitochondrial damage in highly active neurons, which may explain differential sensitivity in disease states.

### MDV and whole organelle mitophagy in neurons

Mitophagy can occur through two distinct methods: production of mitochondrial derived vesicles (MDVs) or whole organelle mitophagy^47,48^. Whole organelle mitophagy is a cargo-selective form of autophagy, the process through which autophagosomes engulf and degrade unwanted cellular components. Damaged mitochondria are targeted to LC3 on the autophagic membrane through LC3 adaptors such as OPTN, NDP52, and TAX1BP1^48–51^. MDV production offers a second method of mitochondria quality control. In this recently discovered process, oxidized proteins accumulate in small, distinct spheres that bud off from mitochondria^23^. These MDVs are then targeted to lysosomes for degradation and recycling^52^. MDV formation is hypothesized to be a baseline or first response to mitochondria stress and damage, as they appear first after cellular treatment with mitochondria damaging agents^53^. Likewise, we noted mitochondrial budding events linked to MDV production in our SBFSEM dataset. Our size analysis of mitophagic vesicles under confocal microscopy further suggests MDVs are the primary method of mitophagy under basal conditions. Only with increased neuronal activity and mitochondria damage does size increase, suggesting the addition of whole organelle mitophagy. However, this will need to be corroborated with electron microscopy analysis, the gold standard for MDV identification.

### Pink1-dependent mitophagy in neurons

In healthy mitochondria, Pink1 is imported to the inner mitochondria membrane, where it undergoes PARL-mediated cleavage and proteasome degradation^54,55^. Upon loss of mitochondria membrane potential, Pink1 is not imported, instead localizing to the outer mitochondria membrane. There, it phosphorylates Parkin, an E3 ubiquitin ligase, that targets the damaged mitochondria to the autophagosome for degradation^9–12^. This mechanism has been engaged previously through the use of mitochondria depolarizing agents such as Antimycin A, CCCP and Killer Red^9,37,56^. Our work, in concert with studies in mammalian systems, suggests that Pink1-mediated mitophagy is not necessary for basal mitophagy. In our model and others, loss of Pink1 or Parkin results in no change in basal mitophagy rates^13,14^. Similarly, many Pink1 knockout animal models show no overt phenotypical changes^14,57^. Likewise, we noticed no phenotypical difference in the axons of our Pink1 knockout animals (Figure S3C). Our findings instead point to a context specific use of Pink1-dependent mitophagy in neurons. In this model, cumulative mitochondria damage and increased activity result in increased mitophagy demand which is met at a mechanistic level by pathways in which Pink1 function is critical. Loss of Pink1 in this context results in wild type levels of mitophagy. This both tells us that Pink1 is required for increased mitophagy and also that basal mitophagy continues in the absence of Pink1.

Together, our work provides pieces of an ever-growing picture of mitophagy regulation and rates in axons of different neuronal populations and suggests a damage and activity specific condition in which Pink1-dependent mitophagy is activated. Taken together, this could help to explain why onset of Parkinson’s disease is relatively late onset in humans and primarily apparent in certain neuronal subtypes.

## Acknowledgements

We thank the members of the Drerup lab for their thoughtful discussions of the work. We thank Randall Massey and the University of Wisconsin School of Medicine and Public Health Electron Microscopy Facility for processing of the serial block face scanning electron microscopy tissue.

## Contributions

C.M.D and S.R.W. designed the research; S.R.W., C.S, E.E.S, M.H., and C.M.D. performed the research; S.R.W, E.E.S, and C.M.D analyzed the data; S.R.W. and C.M.D. wrote the paper.

## Resource Availability Lead contact

Further information and requests for resources and reagents should be directed to and will be fulfilled by the lead contact, Catherine M. Drerup

## Materials availability

Materials are available from the lead contact upon request.

## Data and code availability

The datasets and information required for the data supporting the study are available from the lead contact upon request.

