## Supplemental information for "The combination of elevated neuronal activity and mitochondrial damage induces Pink1-dependent mitophagy in axons"

### Methods and Materials

*Zebrafish husbandry*

All zebrafish (*Danio rerio*) work was done in accordance with the University of Wisconsin- Madison Institutional Animal Care and Use Committee guidelines. Adult zebrafish were kept at 28.5°C in a 14hr/10hr light/dark cycle and embryos were spawned according to established protocols[58]. Embryos and larvae were kept in embryo media (995 µM $MgSO_4$, 154 µM $KH_2PO_4$, 42 µM $Na_2HPO_4$, pH 7.2, 1.3 mM $CaCl_2$, 503 µM KCl, 15 mM NaCl, 714 µM $NaHCO_3$), maintained at 28.5°C and developmentally staged using established methods[59]. Experiments were performed at 5 dpf unless otherwise specified. At this stage, sex is not determined[60]. Genotyping was done after experiments with PCR amplification using primers listed in Table 1.

*Serial Block Face Scanning Electron Microscopy (SFBSEM)*

4 dpf wild type zebrafish larvae were used for pLL axon terminal SBFSEM. To prepare the tissue, tails were cut to isolate the caudal most sensory organs in the tail. Tails were placed into fixative (4% glutaraldehyde in 0.1 M cacodylate buffer) for 1 hour at room temperature and then at 4°C overnight. Following fixation, tissue was washed 5 times for 3 minutes in cold 0.1 M cacodylate buffer. Next, tissue was washed in a solution containing 3% potassium ferrocyanide, 0.3 M cacodylate buffer, 4 mM calcium chloride and 4% aqueous osmium tetroxide for 1 hour on ice. Tissues were then washed with ddH2O 5 times 3 minutes at room temperature before being washed in a 0.22 µm filtered TCH solution (0.1 g thiocarbohydrazide in 10mL ddH2O dissolved in a 60°C oven for 1 hour) for 20 minutes at room temperature. Next, the tissues were rinsed 5 times for 3 minutes in ddH2O, placed in 2% osmium tetroxide for 30 minutes at room temperature followed by 5 additional 3 minute washes in ddH2O and an overnight incubation in 1% uranyl acetate at 4°C. Following overnight incubation, tissues were washed again 5 times for 3 minutes in ddH2O followed by incubation in lead aspartate (0.066 g lead nitrate, 10 mL 0.03 M aspartic acid) solution at 60°C for 30 minutes. After five 3 minute washes in ddH2O, tissues were dehydrated in cold solutions of 20%, 50%,

70%, 90% and 100% acetone for 5 minutes and 10 minutes for the final step. Following dehydration, tissues were processed through a Durcupan series, 25% Durcupan:acetone for 2 hours, 50% Durcupan:acetone overnight, 75% Durcupan:acetone for 2 hours and 100% Durcupan for 45 minutes to 1 hour in 65°C water bath. Resin was then exchanged with fresh 100% Durcupan at 70°C for 20 minutes with 5°C temperature increases every 5 minutes before being placed in molds and polymerizing in a 65°C drying oven for 48 hours.

For mounting, tails were placed on aluminum specimen pins with conductive silver epoxy and sputter coated with pure gold. 3View ultramicrotome was used to remove the top layer of sputter. The neuromast was identified by the characteristic hair cells and support cells protruding from the side. A 2 x 2, 7000 by 7000 pixel montage was acquired, with each pixel equating 5 nm with a 50 nm z-step size.

In the sections, hair cells were identified by their stereocilia at the apex and the presence of the ribbon synapse at the base. Afferent axon terminals were identified by their direct opposition to the ribbon synapses and lack of synaptic vesicles. Efferent axon terminals were identified by the synaptic vesicles present in the terminal synapsing onto the hair cells in a region independent of the presynaptic ribbon. Organelles were identified as previously described [61].

All distance and volume measurements were taken in the ImageJ plugin, TrakEM2. To calculate mitochondria volume, mitochondria were painted with TrakEM2's segmentation function. Areas were taken for each individual mitochondria with z-planes used to calculate volumes. For mitochondria-autophagic vesicle distances, individual distances were taken with the line function from membrane to membrane of each organelle pair. To calculate the number of buds per mitochondria, mitochondria budding events (presence of a vesicle with a constricting neck) were counted per mitochondria and normalized to mitochondria volume calculated above. To all parameters, a $Log_2$ transformation was applied for graphical representation.

*DNA expression plasmids*

DNA expression plasmids used are listed in Table 1. The *mnx:mito-TagRFP-p2a-eGFP* construct was derived using gateway cloning[62]. All other new DNA plasmid constructs were derived using Gibson cloning with the oligonucleotides listed in Table 1. All novel DNA expression plasmids were sequence verified prior to use.

*Live confocal imaging*

Fluorescent indicators were expressed for live imaging using transient transgenesis. For this, 13-25 pg of plasmid DNA was microinjected into zebrafish zygotes as previously described[63]. Plasmids used are listed in STAR Methods Table 1. Zebrafish larvae were identified for expression in pLL or pMN cell bodies using an AxioZoom V.16 Zeiss microscope. For pMNs, we focused only on those between somites 5-22, in the mid-trunk region of the animal[24].

For live imaging, larvae were anesthetized in 0.02% tricaine, mounted in 1.8% low melt agarose in embryo media and imaged with an Olympus FV3000 confocal microscope with a 40x (NA 1.25) silicone oil objective. Optimal interslice interval was used for all imaging. For area measurements, laser power and detector settings were optimized for each animal for precise measurements. For all data analyzed by fluorescence intensity, laser and detector settings were held constant between all animals.

*Stimulation Methods*

pLL sensory neurons were stimulated using water flow. For this, 20-30 larvae were put in a 100 mm petri dish filled half-way with embryo media. The petri dish was rocked on an orbital shaker at 85 rpm for 36 hours from 3.5 dpf to 5 dpf. Larval rocking was maintained until just prior to imaging.

Primary motor neurons were stimulated through bath application of 100 μM N-methyl D-aspartate (NMDA) in embryo media. Control animals were treated with an equal volume of water (NMDA solvent) in embryo media. Groups of 20 were treated in 5 mL of solution in a 6-well plate for 16 hours from 4 to 5 dpf.

*Live imaging analysis*

Mitophagy and autophagy rates were calculated as the ratio of the mCherry only mitochondria or autophagosome area to the total mitochondria or autophagosome area (mCherry+GFP positive). First, a mask was applied to remove all non-mitochondria/autophagosome signal using the mCherry signal. Then, the axon shaft was eliminated to limit analysis to the synaptic regions. Finally, individual GFP and mCherry images were subjected to manual thresholding and area of each measured in ImageJ[64]. mCherry only area was calculated by subtracting the GFP+ area. To confirm measurements, all mitophagy data were subjected to blind interrater analysis.

*pink1 mutant production*

The *pink1* mutant line (*pink1$^{uwd13}$*) was generated using CRISPR/Cas9 mediated mutagenesis as previously described [41]. In brief, gRNAs were generated using CHOPCHOP[65], prioritizing gRNAs in the first exon of the gene. 400 pg of both gRNAs were pooled and injected into zebrafish embryos with 150 ng Cas9 protein (IDT). Embryos were raised to adulthood and outcrossed to wild type (AB) to generate F1 larvae. F1 adults were genotyped using PCR amplification of the targeted region using the primers listed in Table 2 and sequenced to confirm genomic alteration. F1 adults with a 4 bp deletion/2 bp insertion at gRNA 1 site and 48 bp deletion at gRNA 2 site were outcrossed to wild type to generate a stable line.

*Hybridization Chain Reaction RNA Fluorescent In situ Hybridization (HCR RNA FISH)*

HCR RNA FISH was performed as previously described[28,66]. In brief, 4 dpf zebrafish were fixed in 4% paraformaldehyde (PFA) in phosphate-buffered saline (PBS) at room temperature for 2 hours. Fixed larvae were washed in PBS + 0.1% Tween-20 and dehydrated in a methanol series and stored at -20°C. Larvae were rehydrated in a reverse methanol series, treated with proteinase K (10 μg/mL) for 15 minutes and post-fixed for 20 minutes with cold 4% PFA in PBS. Following fixation, larvae were incubated with a manufacturer-supplied hybridization buffer at 37°C for 1 hour and incubated with 4 nM probe in hybridization buffer at 37°C overnight. Following probe incubation, larvae were washed in a manufacturer-supplied wash buffer followed by SSC + 0.1% Tween-20. Next, larvae were incubated in a manufacturer-supplied amplification buffer at room temperature for 30 minutes, during which manufacturer-supplied reporter-labeled fluorescent hairpins were individually heated at 95°C and cooled to room temperature for 30 minutes in the dark. Next, larvae were placed in fresh amplification buffer with 60 nM hairpins and DAPI (1:500). Larvae were then incubated overnight in the dark at room temperature and mounted next day between glass slides with #1.5 coverslips with Fluoromount mounting medium. Larvae were imaged on an Olympus FV3000 confocal microscope with 60x (NA1.42) oil objective. Laser power and detector settings were kept consistent. For imaging, 30 slices at 0.49 μm step size centered around the middle of the spinal cord at the midtrunk were taken for analysis. To quantify signal, spinal cord neuronal cell bodies were isolated and a sum projection was applied through 30 total slices. Mean fluorescence intensity was measured and normalized to wild type.

*TMRE analysis of mitochondrial matrix potential*

Vital dye labeling with tetramethylrhodamine ethyl ester (TMRE) was performed as done previously[31]. To label mitochondria, zebrafish zygotes were microinjected with a DNA plasmid to express eGFP localized to the mitochondrial matrix in motor neurons. At 3 dpf, zebrafish expressing GFP in pMN mitochondria were selected. At 5 dpf, zebrafish larvae were incubated in 25 μM TMRE in embryo media with 0.1% DMSO (dimethyl sulfoxide) in a 12 well-plate for 4 hours in the dark. Experiments were timed such that larvae were in TMRE for precisely 4 hours. Larvae were then washed 3x in embryo media before being mounted in 1.8% low melt agarose and imaged on an Olympus FV3000 confocal microscope with 40x (NA 1.25) silicone oil objective. All laser and detector settings were held constant. For analysis, a mask derived from the eGFP signal was applied to isolate the TMRE signal. Mean fluorescence intensity was calculated from a sum projection through pMN axon terminals.

*Immunohistochemistry of phosphorylated ERK*

Immunolabeling of phosphorylated ERK was used to confirm increased neuronal activity after NMDA treatment. Following 20 hrs of 100 μM NMDA treatment, larvae were fixed in 4% PFA/0.25% Triton X-100 at room temperature for 2 hours. After fixation, larvae underwent an antigen retrieval step where, after being washed briefly in 1x PBS/0.25% Triton X-100 (PBST), they were incubated in 150 mM Tris HCl pH 9.0 first for 5 minutes at room temperature and then for 15 minutes at 70°C. Larvae were then washed briefly with PBST and permeabilized overnight with water at room temperature. Larvae were incubated in block (0.1% Triton X-100, 1% dimethyl sulfoxide, 0.02% sodium azide, 0.5% bovine serum albumin, 5% goat serum) at room temperature for 2 hours, then incubated in rabbit anti-phosphorylated ERK antibody (1:500, Cell Signaling) overnight at 4°C. Following antibody incubation, larvae were washed in PBST then incubated in goat anti-rabbit AlexaFluor 568 (1:1000) and DAPI (1:500) overnight at 4°C. Larvae were then washed with PBST and mounted and imaged as described above for HCR RNA FISH. To quantify signal, spinal cord neuronal cell bodies were isolated and a sum projection applied through 35 total z-steps (0.49 μm interval). Mean fluorescence intensity was measured and normalized to background.

*Statistics and figure assembly*

Image analysis was performed using ImageJ[64]. Statistical analysis was performed using JMP 18 and graphs were made using GraphPad Prism 10. Parametric data was analyzed by ANOVA with Tukey HSD post-hoc contrasts for multiple comparisons. Nonparametric data was analyzed using Wilcoxon/Kruskal-Wallis analysis with multiple comparisons done using the Steel-Dwaas test. Figures were compiled in Adobe Illustrator. Equal and linear adjustments to brightness/contrast were done for all image panels.

**Key resources table**

| REAGENT or RESOURCE | SOURCE | IDENTIFIER |
|---|---|---|
| Antibodies | | |
| Rabbit monoclonal anti Phospho-p44/42 MAPK (Erk1/2) | Cell Signaling | 4370 |
| Chemicals, peptides, and recombinant proteins | | |
| Tetramethylrhodamine ethyl ester | ThermoFisher | T669 |
| N-Methyl-D-Aspartic Acid | MilliporeSigma | M3262 |
| Cas9 Protein | IDT | 10007806 |
| Fluoromount | Sigma-Aldrich | F4680 |
| HiFi DNA Assembly Master Mix | NEB | E2621 |
| Critical commercial assays | | |
| HCR Amplification buffer and RNA-FISH wash buffer | Molecular Instruments, Inc | n/a |
| Zebrafish *pink1* probes | Molecular Instruments, Inc | custom |
| Experimental models: Organisms/strains | | |
| Zebrafish: AB | ZIRC AB | ZDB-GENO-960809-7 |
| Zebrafish: $mitfa^{w2}$ (nacre) | ZIRC | ZL2104 |
| Zebrafish: $actr10^{nl15}$ | Drerup et al.[67] | nl15 |
| Zebrafish: $pink1^{uwd13}$ | This paper | uwd13 |
| Zebrafish: $TgBAC(neurod1{:}eGFP)^{nl1Tg}$ | Obholzer et al.[68] | nl1Tg |
| Zebrafish: $Tg(5kbneurod{:}mRFP\text{-}actr10)^{nl22}$ | Mandal et al.[31] | nl22 |
| Oligonucleotides | | |
| See Table 1 | | |
| Recombinant DNA | | |
| *mnx1:cox8a-cox8a-eGFP-mCherry* | This paper | n/a |
| *5kbneurod1:cox8a-cox8a-eGFP-mCherry* | Wong et al.[28] | n/a |
| *5kbneurod1:Lamp1-TagBFP2* | Liao et al.[69] | n/a |
| *mnx1:mito-TagRFP-p2a-eGFP* | This paper | n/a |
| *mnx1: mCherry-GFP-LC3* | Wisner et al.[41] | n/a |

| | | |
|---|---|---|
| *mnx1: mito-eGFP* | Wong et al.[28] | n/a |
| Software and algorithms | | |
| CHOPCHOP | Labun et al.[65] | https://chopchop.cbu.uib.no/ |
| ImageJ/Fiji | Schindelin et al.[64] | https://imagej.nih.gov/ij/ |
| Prism 10 | Graph Pad | https://www.graphpad.com/features |
| Adobe Creative Cloud | Adobe | https://www.adobe.com/creativecloud.html |
| JMP 18/JMP Student | JMP | https://www.jmp.com/en/home |

**Table of Oligonucleotides**

| | |
|---|---|
| Guide for pink1 g1 | 5’GGCAGGTCGGTTTTCCAGCT**GGG** |
| Guide for pink1 g2 | 5’GCTCCGGACTGGAGCTGCG**CGG** |
| Pink1 fwd | 5’ TCAGTAAAGCATGTTCTCAGCC |
| Pink1 rev | 5’ CTGCTCCTGTTCAATCAGACC |
| Actr10 fwd | 5’ CTGTTTTCGGATGAACTGCCTG |
| Actr10 rev | 5’ ACTTACTTCGTGTAGGCCGC |
| Plasmid generation | |
| *Gata(hb9): cox8a-cox8a-eGFP-mCherry* | |
| Gata fwd | 5’ AAACACAGGCCAGATGGGCCCGAATTGGGTACCGAGCTC |
| Gata rev | 5’ CCCTATAGTGACAAAAGCTGGAGCTCCAC |
| mitoGC fwd | 5’ CAGCTTTTGTCACTATAGGGCGAATTGG |
| mitoGC rev | 5’ TGGATCATCATCGATGGTACTTACTTATACAGCTCGTCC |
| *Gata(hb9): mCherry-GFP-LC3* | |
| Gata fwd | 5’ AAACACAGGCCAGATGGGCCCGAATTGGGTACCGAGCTC |
| Gata rev | 5’ GTGGAGCCTGACAAAAGCTGGAGCTCCAC |
| Mch fwd | 5’ CAGCTTTTGTCAGGCTCCACCATGGTGAG |
| Lc3 rev | 5’ TGGATCATCATCGATGGTACTTACACTGACAATTTCATCCCGAAC |
| *Gata(hb9) mito-eGFP* | |
| Gata fwd | 5’ AAACACAGGCCAGATGGGCCCGAATTGGGTACCGAGCTC |
| Gata rev | 5’ CATGCTAGGCACAAAAGCTGGAGCTCCAC |
| Mito-eGFP fwd | 5’ CAGCTTTTGTGCCTAGCATGTCCGTCCTG |
| Mito-eGFP rev | 5’ TGGATCATCATCGATGGTACCTATAGGGCTGCAGAATCTAGAGGC |

### Supplemental Figure 1

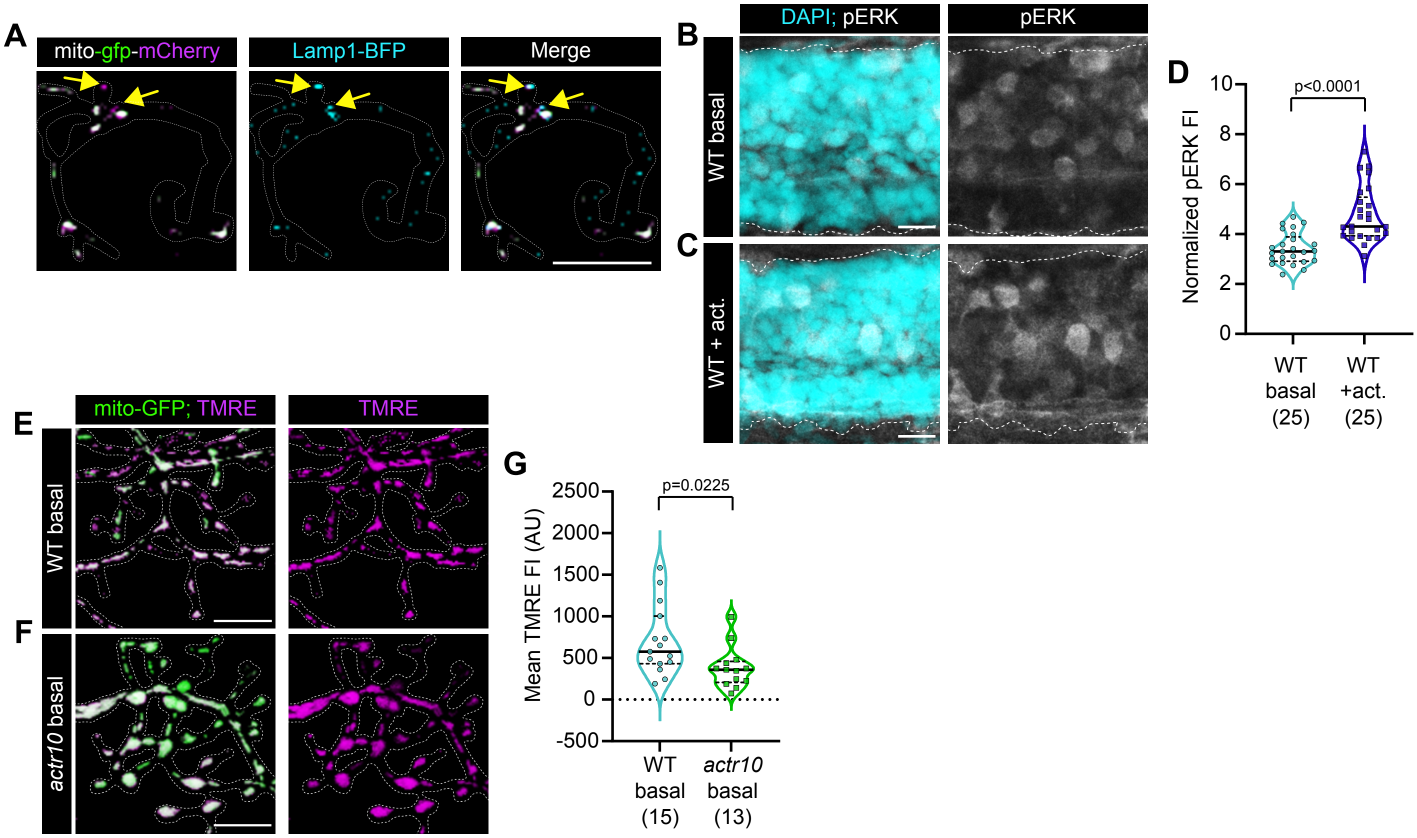

**Supplemental Figure 1 – Mitochondrial matrix potential is reduced in *actr10* mutants pMN axon terminals. Related to Figure 2**

(**A**) Representative image of a pLL axon terminal expressing the mitophagy indicator and the lysosome marker Lamp1-TagBFP2. Arrow indicates colocalization of mCherry only mitochondria (undergoing mitophagy) with a BFP-positive lysosome. (**B**-**D**) Representative images of phosphorylated ERK (pERK) signal (white) in spinal cord (DAPI – cyan) of wild type (WT) zebrafish larvae without (**B**) or with (**C**) increased neuronal activity. (**D**) Quantification of normalized pERK fluorescence intensity (FI; data normalized to non-neuronal signal; Wilcoxon). (**E**-**G**) Representative images of portions of pMN axon terminals of wild type (WT; **E**) and *actr10* mutant (**F**) larval zebrafish expressing mito-eGFP (green) and stained with TMRE (magenta). Scale bars = 10 μm. (**G**) Quantification of mean mitochondrial TMRE fluorescence intensity (FI; ANOVA). Sample size indicated below graphs and represents individual larvae.

### Supplemental Figure 2

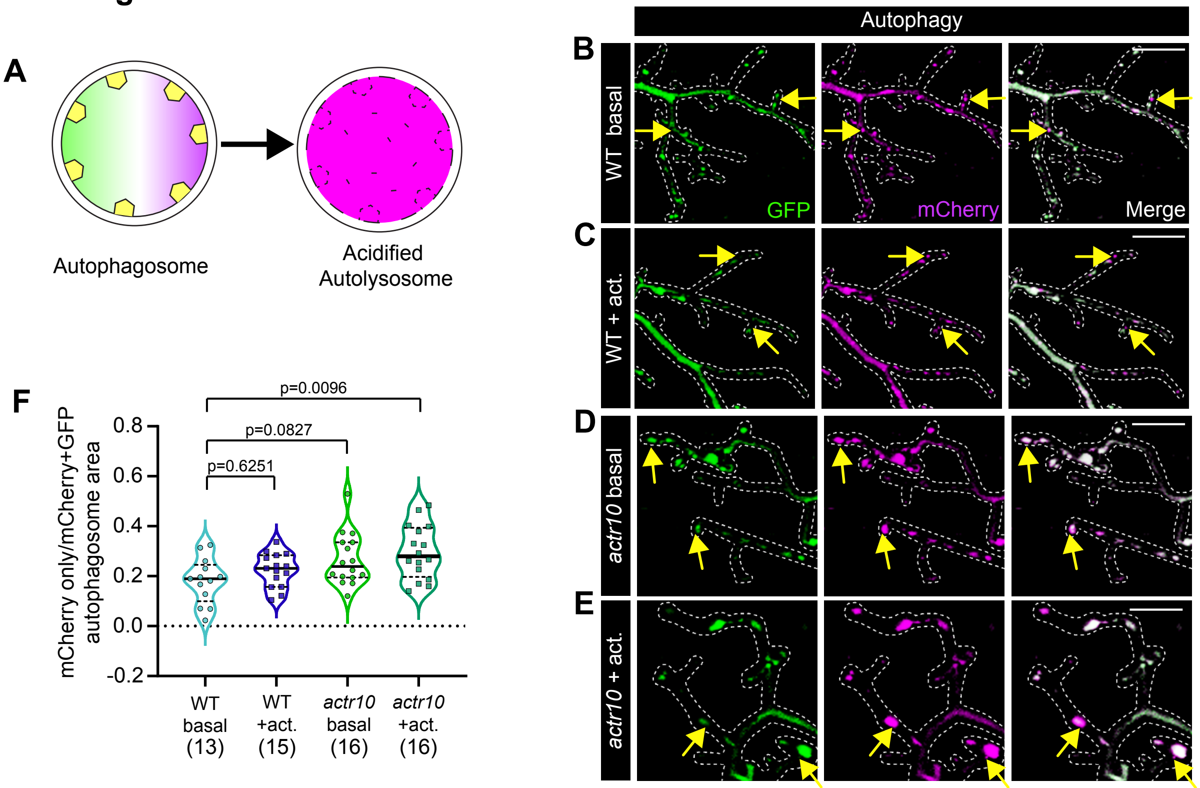

**Supplementary Figure 2 – Autophagy is increased by the combination of mitochondrial damage and neuronal activity. Related to Figure 3**

(**A**) Schematic of the mCherry-GFP-LC3 autophagy reporter. Unacidified autophagosomes are positive for both GFP and mCherry fluorescence whereas autophagosomes that have fused with a lysosome and are undergoing acidification are only mCherry positive. (**B**-**E**) Representative images of pMN axon terminals of wild type (WT; **B**), wild type with increased neuronal activity (WT + act.; **C**), *actr10* mutants (*actr10*; **D**) and *actr10* mutants with increased neuronal activity (*actr10* + act.; **E**) expressing the dual-tagged mCherry-GFP-LC3 plasmid. Arrows indicate acidified autolysosomes. Scale bar = 10 μm (**F**) Quantification of the proportion of acidified autolysosomes to total autophagosome area (ANOVA with Tukey's HSD post-hoc contrasts). Sample size is indicated below graph and represents individual larvae.

### Supplemental Figure 3

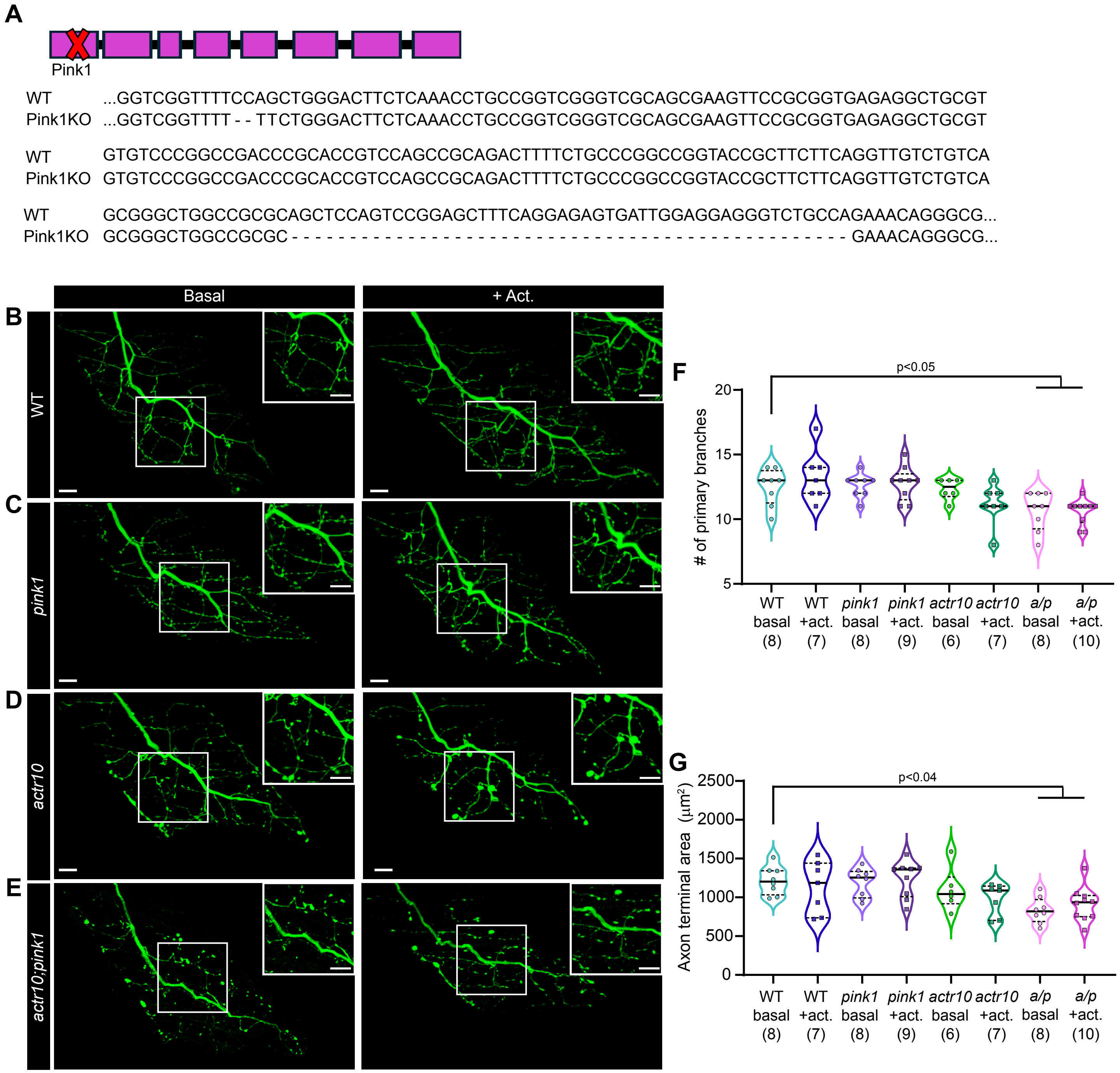

**Supplemental Figure 3 –*actr10;pink1* double mutants have decreased axonal arbors. Related to Figure 4**

(**A**) Schematic of *pink1* exon/intron structure. Red 'X' indicates *pink1* gRNA target sites. Sequence changes shown below. The mutant has a 4 bp deletion/2 bp insertion at one guide site and a 48 bp deletion at the second guide site. (**B**-**E**) Representative images of CaP primary motor neuron axon terminals with or without increased neuronal activity in wild type (WT; **B**), *pink1* mutants (**C**), *actr10* mutants (**D**), and *actr10;pink1* double mutants (**E**). Box indicates region magnified in inset. Scale bars = 10 µm. (**F**) Quantification of the number of primary pMN branches in each condition (*actr10;pink1* as *a/p*, Dunnett's multiple comparisons with wild type basal set as control). (**G**) Quantification of the total axon area starting from first collateral (Dunnett's multiple comparisons with wild type basal set as control). Sample size is indicated below graph and represents individual larvae.
